# Mapping the Mutational Landscape for Streptokinase Binding to Plasminogen

**DOI:** 10.64898/2026.02.04.703780

**Authors:** Srishti Baid, Matthew L Holding, Taylor Laurin, David Seimeiniak, David Ginsburg, Laura M Haynes

## Abstract

Group A Streptococcus (GAS) expresses streptokinase (SK), a critical virulence factor that non-enzymatically activates the host’s plasminogen (PLG), to an active form, PLG_SK_, resulting in the degradation of fibrin clots and subsequent bacterial dissemination. PLG_SK_ formation contrasts with the physiologic activation of PLG to the serine protease plasmin via a proteolytic mechanism. As a potent thrombolytic, SK has been used as a therapeutic to treat heart attacks and strokes. GAS SK is highly specific for human PLG, and sequence variation between SK from different GAS strains has been linked to differences in PLG binding and disease severity. We now report the application of deep mutational scanning (DMS) to map the effects of ∼71% of single amino acid substitutions within SK from Group C Streptococcus, which shares high sequence homology with GAS on its ability to bind human PLG. We first demonstrate that SK expressed as a fusion protein to the p3 coat protein of M13 filamentous phage retains its capacity to bind human PLG. Our subsequent DMS analysis using this phage system identifies regions of SK in which amino acid substitutions are likely to increase or decrease its affinity for PLG. Our findings suggest a complex protein-protein interaction in which long-range protein dynamics influence the conformational activation of PLG to PLG_SK_. These data lay the foundation for linking SK variation between GAS strains to differences in virulence, mapping the determinants of GAS SK’s human specificity, and potentially contributing to the development of improved therapeutics for heart attack and stroke.

## Introduction

Streptokinase (SK) is one of several critical bacterial virulence factor proteins that promote the pathogenesis of Streptococcus species, including Group A (GAS) and Group C (GCS) Streptococcus, by hijacking fibrinolysis (the enzymatic breakdown of fibrin blood clots) and ultimately thwarting the body’s defense mechanism of immobilizing pathogens within thrombi (1-3). GAS is highly specific for humans and causes infections ranging from mild pharyngitis to severe infections, including necrotizing fasciitis (4-6). GCS is an emerging pathogen that can occasionally cross zoological barriers leading to potential human infections (7, 8). SK, which is produced by many pathogenic strains of GAS and GCS, non-proteolytically activates the zymogen plasminogen (PLG) to the potent fibrinolytic enzyme PLG_SK_. Although the general mechanism by which SK activates PLG to PLG_SK_ has been addressed by over three decades of detailed biochemical studies, new tools now enable high-throughput analysis of the biochemical functions for individual amino acids within SK. These approaches should also provide an improved understanding of sequence diversity among SK from different GAS and GCS isolates (9).

Under physiologic conditions within the vascular system, PLG is proteolytically activated to the serine protease plasmin by either tissue- or urokinase-type plasminogen activator (tPA or uPA, respectively) via cleavage of the Arg^561^-Val^562^ peptide bond (10). In contrast, SK non-proteolytically activates PLG to PLG_SK_ by first binding to PLG and then inducing a conformational change by inserting SK Ile^1^ into the activation pocket within PLG (11). PLG_SK_ also has a unique activity profile relative to plasmin, including an enhanced substrate specificity for PLG to generate plasmin (12). Furthermore, SK has a higher affinity for plasmin than for PLG, with SK from some strains of GAS protecting plasmin from inhibition by its primary inhibitor α_2_-antiplasmin(9).

SK from the GCS, *Streptococcus dysgalactiae* subsp. *equisimilis* (SDSE) H46A strain (SK_H46A_) shares high sequence homology with SK from select GAS strains (∼90%) and is one of the best characterized SK variants due to its use as a clinical thrombolytic agent to resolve thrombi associated with acute strokes and myocardial infarctions (11, 13-15). While tPA is currently the preferred thrombolytic treatment in high-income countries, SK_H46A_ is still a first-line therapeutic in middle- and lower-income countries due to its lower production costs (16), despite its reduced efficacy compared to tPA (17).

The SK gene consists of 1398 nucleotides, corresponding to 414 amino acids. The SK protein is composed of three distinct structural domains that each appear to be required for PLG activation (18, 19), with each containing unique functionalities (18, 20-22). The α domain (residues 1-150) is the major domain facilitating substrate recognition and the conversion of PLG to an active conformation; the β domain (residues 151-287) enhances binding affinity and substrate processing during binding; and lastly, the γ domain (residues 288-414) stabilizes the SK-PLG complex and enhances proteolytic activity (23, 24). While the general functions of each of these domains have been identified, we do not yet know the role of individual amino acids in supporting these functions and ultimately the contribution of each amino acid residue with respect to SK’s affinity for and activation of PLG.

We now report deep mutational scanning (DMS) using a high-throughput phage display system (25-28) to simultaneously analyze the role of approximately 70% of all possible single amino acid substitutions in SK with respect to its ability to bind human PLG. Our results provide high-resolution insight into which regions/domains of SK are tolerant of amino acid substitutions with respect to PLG binding, building a platform for the characterization of SK variants from emerging strains of GAS and GCS, as well as the development of potential novel SK therapeutics.

## Results and Discussion

### Phage displayed SK_H46A_ selectively binds, but does not activate human PLG

To demonstrate that SK_H46A_ is capable of specifically binding to PLG when displayed on M13 filamentous phage, we cloned wild-type SK_H46A_ with an N-terminal His_6_-tag and TEV protease (TEVp) cleavage site (**Figure 1A**) into the modified pAY-FE plasmid vector (Genebank #MW464120) (27, 29). We then expressed this phage displayed construct along with a control phage expressing the A3 domain of von Willebrand factor (VWF-A3), as previously described (25, 26). A mixture of SK_H46A_ and excess VWF-A3 phage selected for binding to biotinylated Glu-PLG; (**Figure 1B**), demonstrated strong enrichment for SK_H46A_. – with ∼3% of the unselected phage identified by sequencing as SK_H46A_, compared to ∼90% of the selected phage. These results confirm that phage displayed SK_H46A_ can selectively bind to biotinylated PLG.

**Figure 1:**
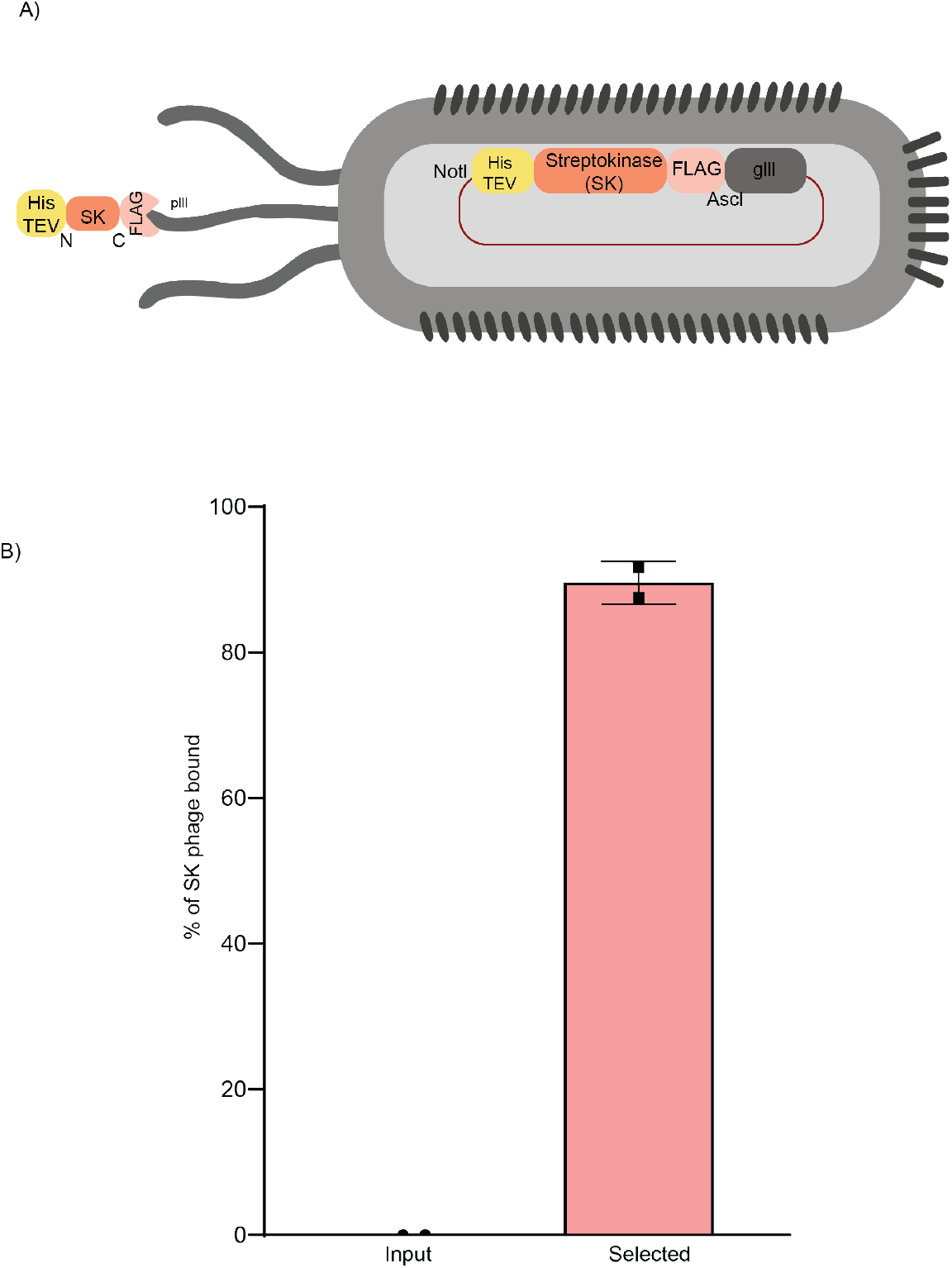
Phage display of functional recombinant SK. A) SK was displayed on M13 filamentous phage by fusion with truncated pIII protein with an N-terminal His-TEV and C-terminal FLAG tag. The NotI and AscI restriction sites flank the His-TEV and the FLAG sequences, respectively, to facilitate cloning of the recombinant SK DNA into the phagemid pAY-FE plasmid vector, as previously described (25, 26). B) Phage displayed SK was incubated with an excess of VWF A3 domain phage (Input), selected for binding to biotinylated plasminogen (Selected). Individual colonies (36 colonies for each condition, for a total of 2 experiments) of phage were genotyped by PCR with flanking primers (Supplementary Table 1) and run on agarose gel and are represented as a percentage of total clones screened for each condition, as described in *Materials and Methods*.

The N-terminal Ile residue (Ile^1^) in SK is known to be critical for its activation of PLG to PLG_SK_ (21); however, our construct design masks this function of Ile^1^ by the presence of the N-terminal His_6_ tag followed by a TEVp cleavage site (see *Materials and Methods*). Previous studies suggest that SK with a blocked or absent Ile^1^ may still exhibit residual PLG activation activity((30)). To detect any potential residual SK activity for our phage displayed fusion protein, we measured PLG activation in an amidolytic assay (**Supplementary Figure 1**) with the fluorogenic substrate (z-Gly-Gly-Arg-7-amido-4-methylcoumarin (AMC)). No residual PLG_SK_ activity was detected under the tested experimental conditions.

### Characterization of the SK_H46A_ variant library

Our error-prone PCR-generated SK_H46A_ variant library exhibited a depth of ∼10^7^ independent clones, with DNA sequence analysis of 24 randomly selected phage demonstrating an average of ∼2 amino acid substitutions per clone as determined by Sanger sequencing (**Supplementary Figure 2**). High-throughput sequencing (HTS) of the variant library revealed that it contained 5,603 of 7,866 possible single amino acid substitutions (71%). As expected, the coverage frequency for individual amino acid substitutions correlates with their number of codons in the genetic code, with significant underrepresentation of amino acid residues encoded by a single codon (**Supplementary Figure 3**).

### DMS of SK_H46A_ binding to PLG

We next assessed the ability of variants in the SK_H46A_ library to bind PLG, using the approach depicted schematically in **Figure 2A**. Briefly, the SK_H46A_ variant library was incubated with 100 nM biotinylated PLG, and phage displayed SK_H46A_ variants in complex with biotinylated PLG were selected with streptavidin resin. The segment of selected phage DNA encoding SK_H46A_ was amplified and sequenced in twelve 150 bp amplicons (**Figure 3A**) (27). Principal component analysis (PCA) demonstrates reproducibility in our selection assays (**Figure 2B**) with clear clustering of replicates for the input phage library, biotinylated PLG selected phage, and non-specific binding conditions (no PLG control). To identify regions of SK_H46A_ in which amino acid substitutions exhibited the largest effects on PLG binding, we also performed individual PCAs for each amplicon (**Figure 3B**) (27). The non-specific binding control (without added PLG, designated as SK-PLG) was consistently separated from the reaction mixture (SK+PLG), and Input (-PLG) along the PC1 in each PCA. However, the explanatory power of PC1 varied considerably among the twelve amplicons—most markedly (>80% variation explained by PC1) for amplicons 1 (residues 1-23), 2 (residues 24-56), 4 (residues 94-130), 5 (residues 131-164), and 8 (residues 234-268), suggesting that amino acid substitutions in these regions are more frequent effectors of PLG binding.

**Figure 2:**
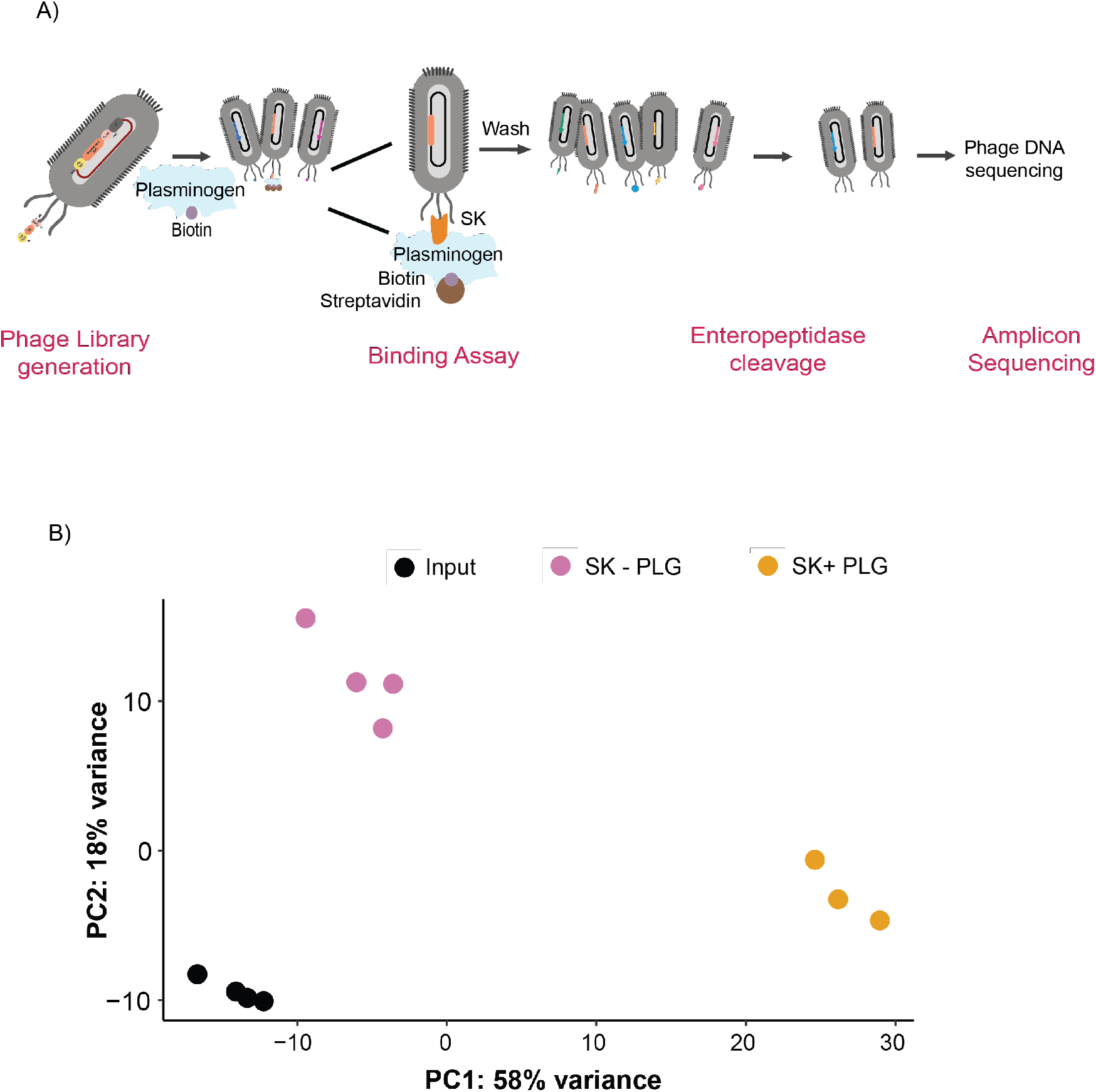
Selection of variants in the phage displayed SK library by affinity for biotinylated PLG exhibits distinct clustering. A) Schematic of the procedure to measure PLG binding by the SK phage library. Phage were isolated and incubated with biotinylated PLG, collected by immunoprecipitation with streptavidin beads, followed by High throughput DNA sequencing (HTS). B) PCA plot shows the correlation of data points among the control reaction (SK-PLG) (pink), the input (black) and the experimental reaction (SK+PLG) (yellow). Replicates (n=4 for inputs and control and n=3 for the experimental samples) for each reaction cluster together, with high variance between the experimental sets.

**Figure 3:**
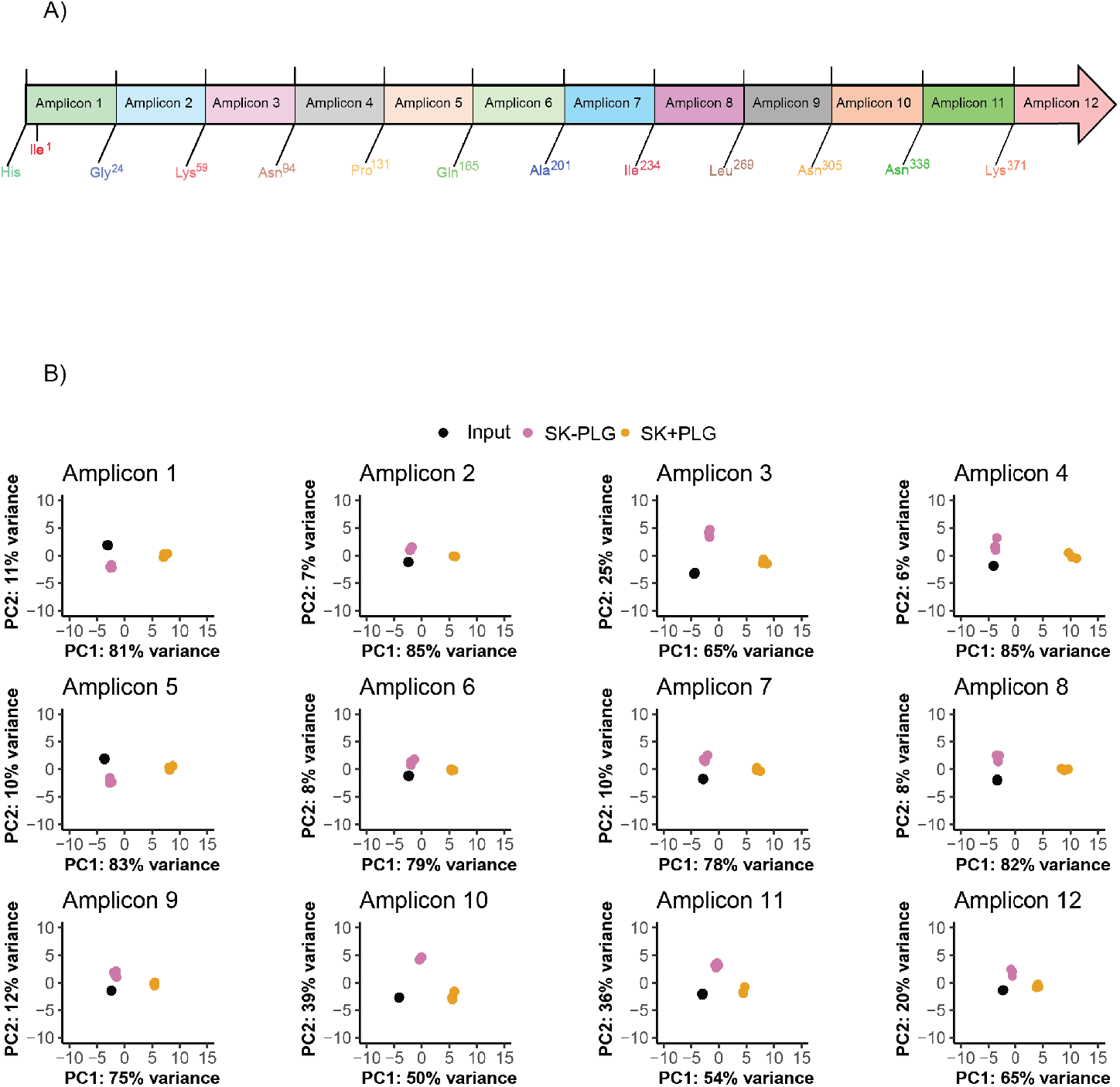
Specific amplicons in SK correspond to regions critical for PLG binding. A) Schematic representation of the SK sequence (1398 bp) showing each amplicon along with the included amino acids. Each amplicon was approximately 150bp in length. B) PCA plots show comparison (log_2_-fold enrichment scores with respect to binding) among the 12 different amplicons across Input (black, n=4), SK-PLG (pink, n=4) control and SK+PLG (orange, n=3) reaction sets to show region specific effects.

Of the amino acid substitutions present in the variant library, 1,670 (∼30%) exhibited significant enrichment (binding score > 0; n = 793; base mean ≥ 100, FDR-adjusted p < 0.1) or depletion (binding score < 0; n = 877; base mean ≥ 100, FDR-adjusted P < 0.1) as shown in **Figure 4A** (and **Supplementary Figure 4**). We interpret amino acid substitutions in SK_H46A_ with binding scores greater than zero as retaining or increasing affinity for biotinylated PLG relative to wild type (WT) SK_H46A_. Those with binding scores less than zero, we interpret as having decreased affinity for biotinylated PLG. Consistent with their propensity to disrupt secondary structures, Pro (74%) and Gly (58%) substitutions were particularly likely to be associated with negative binding scores.

**Figure 4:**
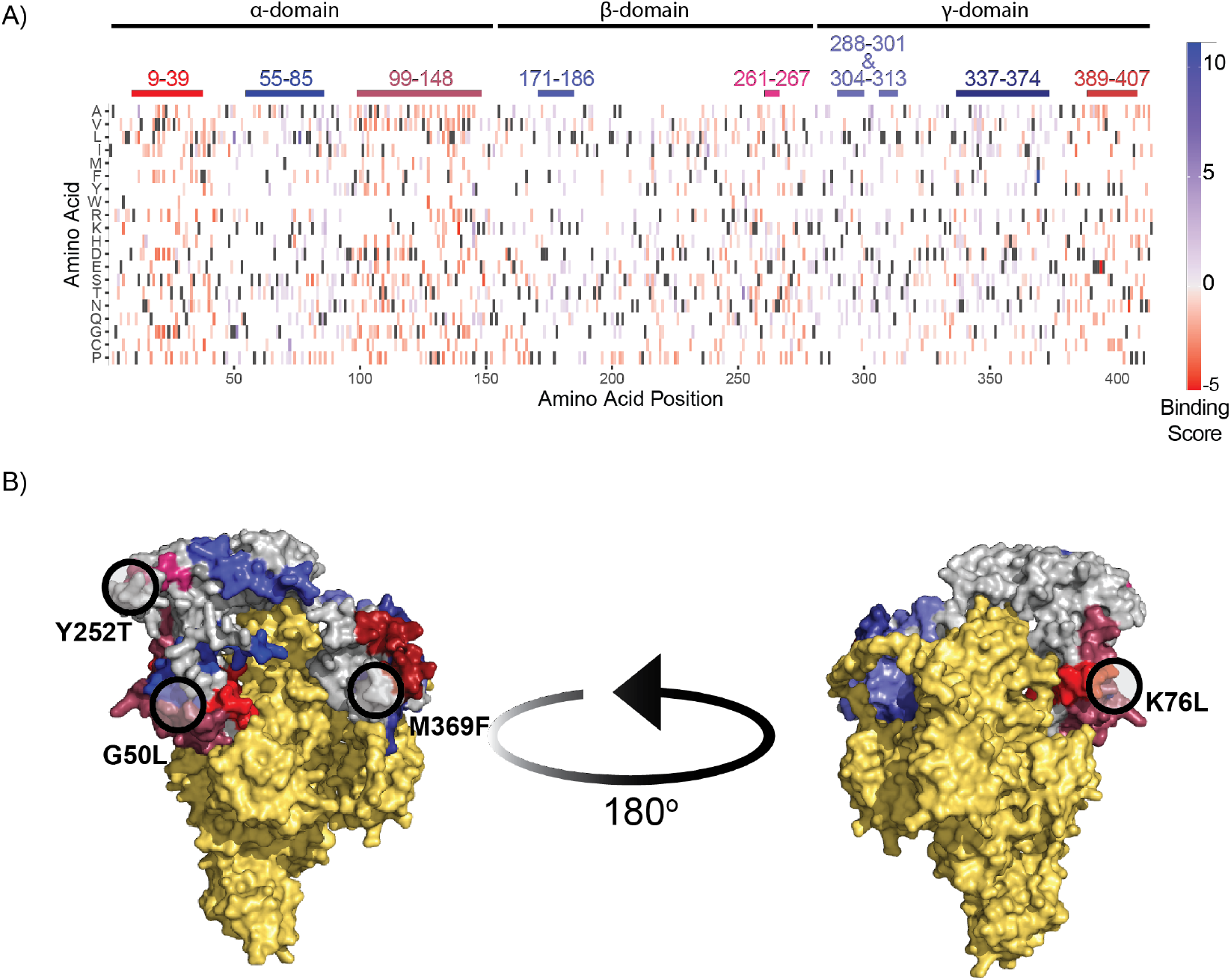
Heat map of DMS analysis and structure alignment of SK and PLG interaction show amino acids critical for binding. A) The consolidated heat map at each position of the SK amino acid sequence indicating those mutations that result in either loss of PLG binding (red) or maintained/enhanced binding (blue). Amino acid positions are indicated on the x-axis, with the y-axis showing the amino acid substitution at each position. The charcoal grey indicates residues and white indicates lack of coverage at that position. B) An AlphaFold (48) model of the SK PLG complex is shown, with SK in grey and PLG in yellow. Selected domains of SK are highlighted in the colors corresponding to the colored segments at the top of panel A – residues 9-39 (red),55-85 (blue), 99-148 (maroon), 171-186 (purple), 261-267 (pink), 288-301 and 302-313 (purple) 337-374 (blue), 389-407 (red). The figure is rotated 180° on the left. The residues that were expressed and validated for activation experiments (**Figure 6**) are highlighted with black circles.

To further characterize segments of SK_H46A_ in which amino acid substitutions are particularly likely or unlikely to disrupt SK binding to PLG, we calculated a *mutational acceptance score* (27, 31). We then fit the positional binding score to a LOWESS regression to determine regions in the SK_H46A_ primary sequence in which amino acid substitutions were likely to maintain, increase, or decrease the affinity of SK_H46A_ for biotinylated PLG (**Figure 4** and **Supplementary Figure 5**). This analysis identifies five regions in SK_H46A_ (75^th^ percentile or greater) in which amino acid substitutions are particularly likely to maintain or increase SK_H46A_’s affinity for PLG (**Figure 4A**; residues 55-85, 171-186, 288-301, 304-313, and 337-374) and four regions (25^th^ percentile or less) in which amino acid substitutions decrease SK_H46A_’s affinity for PLG (**Figure 4A**; residues 9-39, 99-148, 261-267, and 389-407). These regions correspond to segments of SK in which amino acid changes are known to result in a decreased ability for SK to bind and activate PLG to PLG_SK,_ including the N-terminal alpha-domain (12, 32, 33), the 250-loop (20, 34, 35), and the C-terminal Lys^414^ (36).

One limitation of our current assay is that it cannot distinguish between decreased affinity due to protein misfolding or to more subtle structural changes that decrease the binding affinity of a well-folded protein. To distinguish between these two possibilities, we calculated ΔΔG values for amino acid substitution in SK_H46A_ using two computational protein stability predictors, EvoEF2 (37) and FoldX (38) whichare moderately correlated with each other (R^2^ = 0.2930, p < <0.0001; **Supplementary Figure 6**). Amino acid substitutions resulting in decreased thermodynamic stability and protein misfolding should exhibit more positive ΔΔG values, while those that increase thermodynamic stability should exhibit more negative ΔΔG values, suggesting the presence of properly folded proteins, as most proteins canonically fold to their lowest energy state. As anticipated, both EvoEF and FoldX (**Figure 5**), predict that SK_H46A_ variants that bind PLG (binding score > 0) have more negative ΔΔG values than those that exhibit a reduction in binding affinity (binding score < 0). Of note, there is a population of SK_H46A_ variants with binding scores less than zero but with ΔΔG values similar to those variants with binding scores greater than zero, potentially representing a subpopulation of variants that are properly folded yet still exhibit loss of binding.

**Figure 5:**
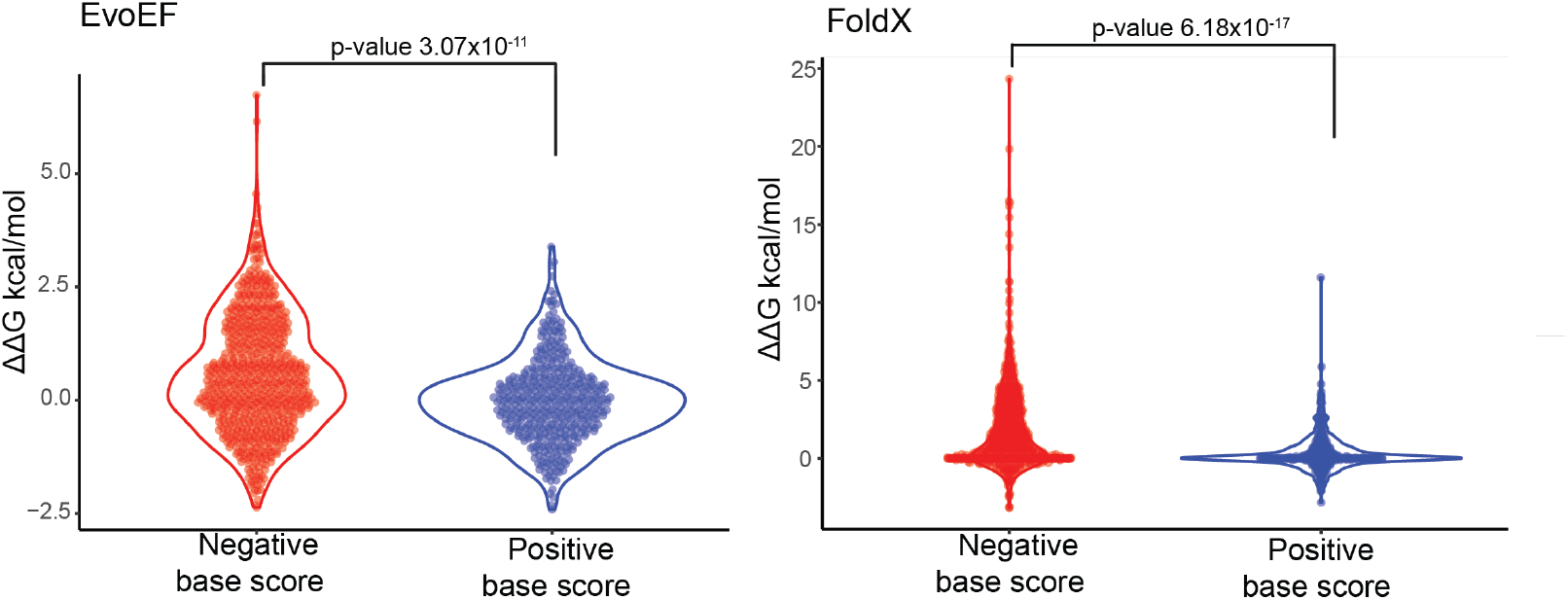
Stability predictions (ΔΔG values) for mutants, using 2 models. EvoEF and FoldX predict that SKH46A variants with higher PLG binding (n=754; binding score > 0) have more negative ΔΔG values than those with reduced binding (n=448; binding score < 0). Select SKH46A variants exhibit low binding scores but ΔΔG values similar to strong binders, suggesting these are properly folded proteins with impaired binding. Amino acid substitutions that decrease thermodynamic stability and cause misfolding show higher (more positive) ΔΔG values, while those that stabilize the protein have lower (more negative) ΔΔG values, consistent with proper folding.

### Characterization of amino acid substitutions that maintain or increase SK-affinity for PLG

Our DMS analysis also identified amino acid substitutions in SK_H46A_ that appear to facilitate SK binding to PLG (**Figure 4**). The amino acid substitutions with the highest binding scores include M369F (11.1), K76L (8.4), G50L (6.1), and Y252T (5.3). (**Figure 4B**; **Supplementary Figure 5**). To validate these findings and explore the effect of these substitutions on SK activation of PLG to PLG_SK_, we recombinantly expressed each of the above individual variants in an *E. coli* expression system, with the same N-terminal His_6_-tag and TEV protease cleavage sequence as in our phage displayed construct. The SK_H46A_ variants were purified, and the His_6_-tag removed by treatment with TEV protease (see *Materials and Methods)*. The recombinant SK_H46A_ variants were then used in a modified plasma-based halo/HoFF assay to simultaneously measure PLG binding and activation to PLG_SK_ (39, 40). No significant differences were observed in the maximum amounts of plasmin generated (**Figure 6A**) nor the maximum rates of fibrinolysis (**Figure 6B**) between the TEV protease treated SK_H46A_ WT and any of the variants, suggesting that amino acid substitutions in SK_H46A_ that increase its affinity for PLG do not necessarily improve nor impair its ability to induce PLG fibrinolytic activity. Furthermore, none of these variants are located at the predicted SK_H46A_-PLG protein-protein interface (41), suggesting that these amino acid substitutions may impact binding affinity via long-range interactions.

**Figure 6:**
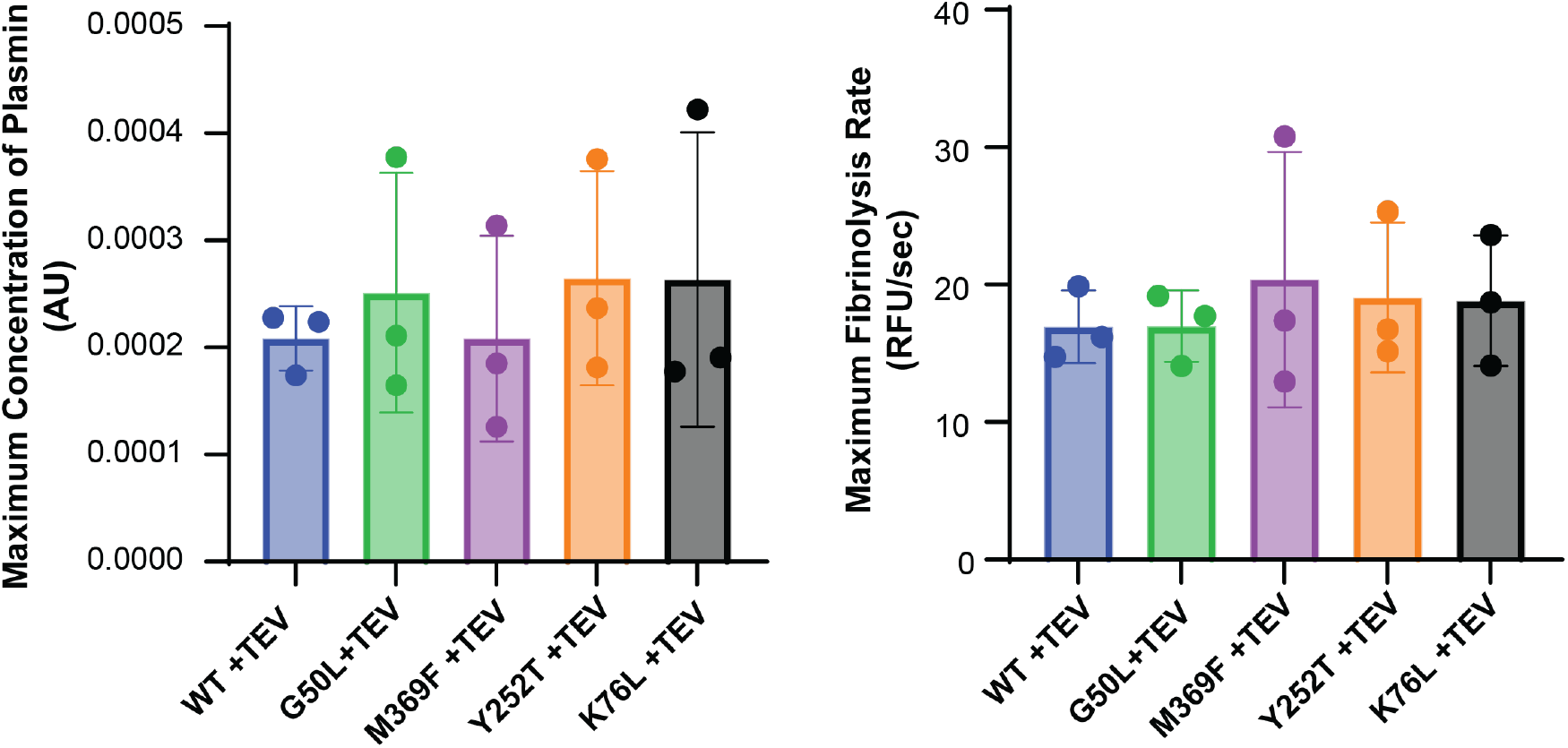
Assessment of select mutants show similar levels of plasmin activation and fibrinogen breakdown as compared to wildtype. The select SK variants (G50L, M369F, Y252T and K76L) were expressed, treated with TEV protease, to free the Ile^1^ and checked for activity by performing a modified Halo assay (described in Materials and Methods). A) The absorbance (λ_abs_ = 405nm) was measured for plasmin activity using plasminogen chromogenic substrate S-2551 and B) Fluorescence (λ_ex_ = 488 nm and λ_em_ = 530 nm) was measured for clot fibrinolysis using fibrinogen simultaneously. Statistical analysis using ANOVA suggested no significant difference in activity between wildtypes and mutants for both experiments.

To further examine the role that long-range interactions, play in the binding of SK_H46A_ to PLG, we evaluated the propensity of an amino acid position to accept amino acid substitutions as a function of its relative solvent accessibility (RSA; **Figure 7**). When comparing orthologous sequences among different species, amino acid variation is most frequent at surface exposed residues (42). One might thus anticipate that amino acid positions in SK_H46A_ that are most accepting of substitutions will generally be those that are surface exposed. However, our DMS screen data suggest that single amino acid substitutions that most improve SK_H46A_’s affinity for PLG are not located at the protein-protein interface. Indeed, both the Met^369^ and Lys^76^ variants characterized above are buried residues, with RSA scores of zero. However, we observe a weak yet significant positive correlation between the positional binding scores and RSA (slope = 1.1, R^2^ = 0.1, p = 1.12×10^-11^), with core residues (RSA < 0.2) generally less accepting of amino acid substitutions, indicating their structural importance in SK, though some core residues in SK are accepting of amino acid substitutions. These deviations from expectations align with recent models suggesting that amino acid substitutions at a protein’s core that do not cause gross protein misfolding can modify a protein’s binding interactions (43, 44).

**Figure 7:**
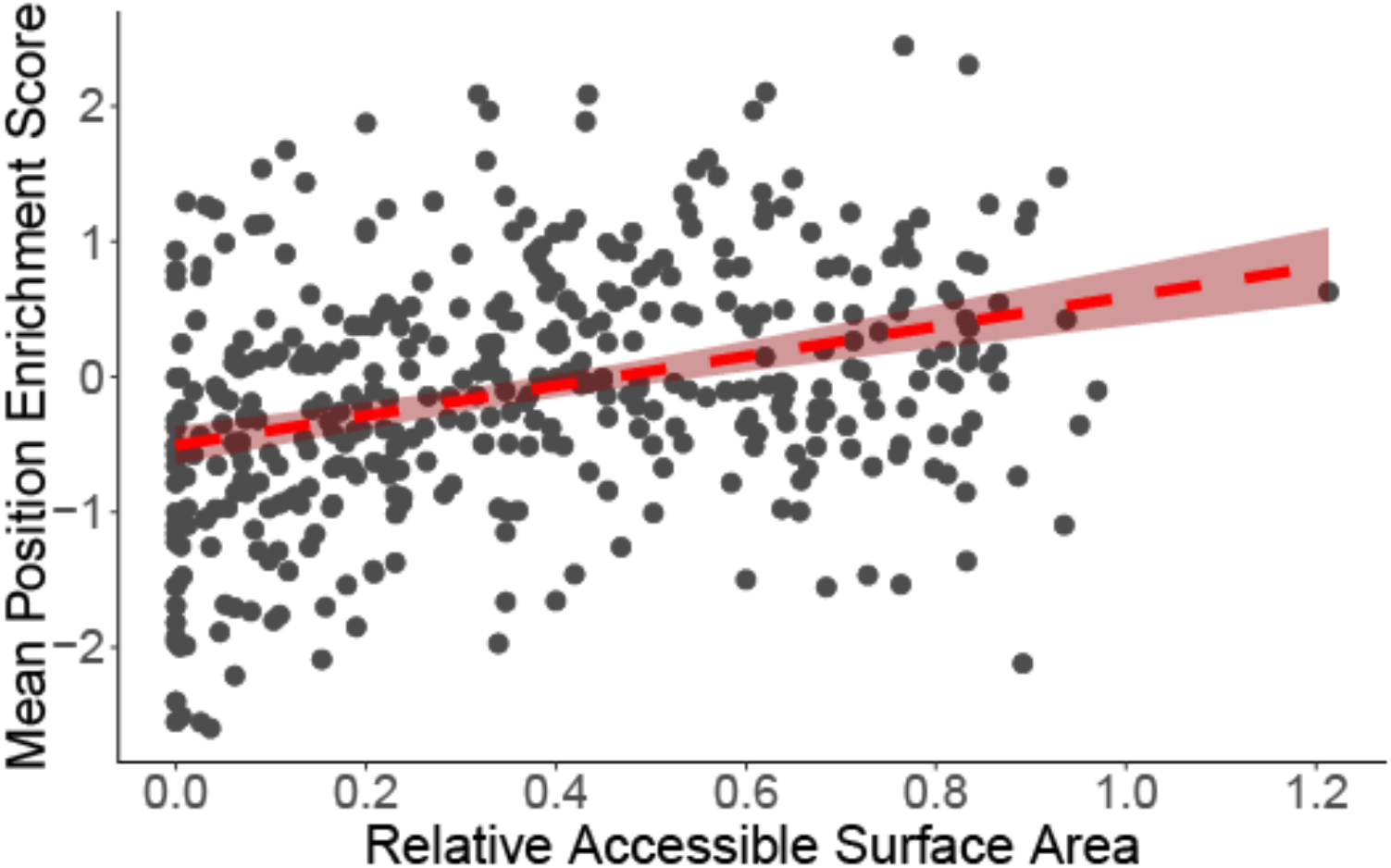
Amino acid substitutions in surface-exposed SK residues are more likely to enhance PLG binding. The mean positional enrichment scores (log_2_-fold change) for each site in SK (from **Figure 4**) are plotted against the relative solvent-accessible surface area (RSA) as predicted by the AlphaFold2 SK structure (see Methods and Figure 4D). Residues with RSA values below 0.2 are typically buried within the protein core, while higher RSA values denote greater surface exposure. Elevated mean enrichment scores correspond to positions more tolerant of substitutions, whereas more negative values indicate lower tolerance for amino acid changes. The red line depicts the best-fit linear regression, with a 95% confidence interval, yielding a slope of 1.1, R^2^ of 0.1, and p-value of 1.12 × 10^− 11^

## Conclusions

Our study complements a growing number of reports using DMS to characterize mediators of infectious disease (45), while laying the foundation for studying bacterial evolution and pathogenesis. Our results suggest that amino acids beyond the site of direct interaction with PLG affect SK-PLG binding through long-range interactions. Finally, this work establishes a valuable foundation for future research into bacterial pathogenicity, particularly at the host-pathogen interface. By mapping the effects of specific mutations within SK on its interaction with human PLG, we provide a resource that will enable deeper investigations into the molecular mechanisms underlying bacterial virulence and adaptation. Furthermore, the insights gained from this study have potential translational applications, as they may guide the development of new and improved thrombolytic agents that leverage engineered SK variants with tailored activity or specificity.

## Materials and Methods

### SK_H46A_ cloning and variant library generation

The coding sequence for SK_H46A_ from SDSE (Genbank ID: CAA51351.1) was synthesized by Twist Bioscience (South San Francisco, CA) and was cloned into a modified pAY-FE plasmid (GenBank ID: MW464120) that carries an ampicillin resistance marker, in which the amber stop codon (*TAG*) immediately preceding the SK-gIIII fusion protein was mutated to a glutamine residue (CAG, Gln), between the AscI (Product# R0558S, New England Biolabs, Ipswich, MA,USA) and NotI (Product # R3189S, New England Biolabs) restriction sites (Supplementary table 1) using T4 ligase (Product #M1794, Promega, Fitchburg, WI, USA) (26, 27). At the N-terminus of SK, a His_6_-tag followed by a TEV protease cleavage site (12, 29) was introduced, while the C-terminus harbored a FLAG-tag prior to the p3 coat protein. The final construct (**Figure 1A**) was transformed into chemically competent XL-1 Blue cells (Agilent Technologies, Santa Clara, CA, USA).

### SK variant library preparation

The SK variant library was generated by error-prone PCR using the GeneMorph II Random Mutagenesis Kit (Product# 200550, Agilent Technologies). Primers used for PCR mutagenesis (**Supplementary table 1**) maintained the AscI and NotI restriction sites for ligation of the restriction-digested insert into the modified pAY-FE vector described above. The PCR products were further amplified with GoTaq (Product# M7123, Promega) using the same primers described above and gel purified using the Qiagen Gel Purification Kit (Product # 28704, Qiagen, Venlo, Netherlands). The resulting library was restriction digested with AscI and NotI and ligated using T4 ligase into the modified pAY-FE plasmid. Following ligation, the library was transformed into electrocompetent XL-1 Blue MRF’ *E. coli* per manufacturer’s instructions. The depth of the library was determined by titering the number of ampicillin-resistant colonies on LB agar (Product #22700025, Fisher Scientific, Waltham, MA, USA) plates supplemented with ampicillin (100 mg/mL) and 2% glucose. Mutation frequency was assessed by Sanger sequencing of randomly selected individual colonies (n=24), and the range of the number of mutations was plotted.

### Expression of phage displayed SK

XL-1 blue cells harboring plasmids for the SK- or VWF-A3-pIII fusion proteins were grown in LB media (Invitrogen) supplemented with 2% glucose (LBG; 100 mL) containing 100 mg/mL ampicillin to mid-log phase at 37 °C and then inoculated with M13K07 Helper Phage (10^14^ CFU/mL; Cytiva, Marlborough, MA, USA) expanded for 1 h at 37 °C, and quantified following manufacturer’s instructions. *E. coli* were pelleted by centrifugation (4,500xg for 10 min at 4 °C) and resuspended in 2xYT Broth (Novagen, Product # 71755-4, Millipore Sigma, Burlington, MA, USA) supplemented with 100 mg/mL ampicillin, 30 mg/mL kanamycin, and 0.4 mM IPTG, and grown for 2 hours at 37°C. *E. coli* were pelleted by centrifugation successively at 4,200xg, followed by 4,500xg for 10 min at 4°C. Phage in the supernatant were precipitated with PEG-8000 (2.5% w/v) and NaCl (500 mM) overnight at 4°C, pelleted by centrifugation (20,000xg) and resuspended in 1X TBS (0.05M Tris Base, 0.15M NaCl; pH 7.4).

### Plasminogen biotinylation

Human Glu-PLG (HCPG-0130; Prolytix; Essex Junction, VT, USA) was biotinylated using NHS biotin (Product # 21925, ThermoFisher, Waltham, MA, USA) per manufacturer’s directions. Free biotin was removed by dialysis against 1X HBS (20 mM HEPES containing 150 mM NaCl; pH 7.4). Protein integrity and biotinylation were confirmed by gel electrophoresis (Invitrogen Wedge Well Novex 4 to 20% TrisGlycine, Product# XP04200BOX, Invitrogen) followed by Coomassie staining. For western blots, proteins were transferred to a nitrocellulose membrane (iBlot™ 2 Transfer Stacks, nitrocellulose, mini, Product# IB23002 XP04200BOX, Invitrogen), which was subsequently blocked with 5% bovine serum albumin (BSA) in TBS containing 0.1% Tween-20 (TBST). The blot was probed with streptavidin conjugated to HRP (1:1000) (Product#N100, ThermoFisher) prepared in TBST containing 5% BSA, washed with TBST, developed with SuperSignal™ West Femto Maximum Sensitivity Substrate (Product# PI34095, ThermoFisher), and imaged.

### Screening of SK variant affinity for PLG

Phage displayed SK was screened for PLG binding using an approach adapted from our previous studies of plasminogen activator inhibitor-1 (25-27, 31, 46). Assay conditions were optimized by confirming that phage displayed SK could be selectively enriched for their PLG affinity compared to excess negative control-phage displayed VWF-A3 (1SK: 9VWF-A3 volume:volume mixture) (25). Once the expression was confirmed, our assay was performed using the SK error-prone PCR library. The reaction was initiated by incubating 50ul of concentrated SK phage (∼10^13^) with biotinylated 100nM Glu-PLG in 1×HBS buffer supplemented with 5% bovine serum albumin (BSA) and incubated at room temperature for 30 minutes on a rotator. Subsequently, the mixture was added to magnetic streptavidin beads (Product # S1420S, New England Biolabs) and incubated overnight at 4 °C with continuous rotation. The bead-bound complexes were then washed with 1×HBS buffer containing BSA. Enteropeptidase (EK) (Product #P8070L, New England Biolabs), which cleaves C-terminal to the FLAG tag, was then added in 50 uL of EK buffer (200 mM Tris, 50 mM NaCl, 2 mM CaCl_2_, pH 8) and incubated overnight at 4 °C. In parallel, XL1 cells were inoculated for subsequent phage titer determination. After incubation, the supernatant was separated from the magnetic beads, and the beads were washed with EK buffer to achieve a final volume of 100 µL. The recovered supernatant containing the phage particles was serially diluted, incubated with XL1 cells grown to an OD600 of 0.4–0.6 for 1h at 37 °C, and 100 µL aliquots were plated onto agar plates supplemented with 2% glucose and ampicillin (100 μg/mL). Colony forming units were enumerated to assess phage enrichment relative to control samples. In competition assays (n=2) with 9 fold excess VWF (volume:volume), random colonies (n= 24) were genotyped by PCR as previously described (25) using primers that anneal to region of the plasmid common to both SK and the VWF-A3 constructs (**Supplementary Table 1**; SK cloning seq check FP, SK cloning seq check RP) and the results were plotted as percentages on Graphpad.

### High-throughput DNA sequencing

Libraries for sequencing were generated as previously described (25). Briefly, primers were designed to generate twelve partially overlapping 150 bp amplicons from pAY-FE error-prone SK via PCR (**Supplementary Table 1**), with each overlapping sequence assessed in only one designated amplicon (**Figure 3A**). PCR products were purified from agarose gels and combined so that the pooled DNA (100 ng) represented equal contributions from each of the twelve amplicons. The mixed amplicons were end-repaired and dA-tailed using the NEBNext Ultra End Repair/dA-tail kit (New England Biolabs), followed by ligation to NextFlex barcoded adapters (Bioo Scientific, Austin, TX, USA) utilizing the NEBNext Ultra Ligation kit (New England Biolabs). Ampure beads (Beckman Coulter, Indianapolis, IN, USA) were used for purification in accordance with the manufacturer’s protocol. HTS (2×150bp paired-end reads) was performed on the Illumina platform (Illumina, San Diego, CA, USA) by MedGenome (Foster City, California, USA). Sequence data analysis was performed with DESeq2 (47) (10% FDR cut off), evaluating mutation frequency at each nucleotide position independently of other mutations present in the SK coding sequence.

### Amidolytic assays

To check for background PLG activation during our SK phage binding protocol, we measured PLG activation at the incubation points during the course of the binding experiment (n=2). We added 100 µL of the reaction mix to the substrate after 30 minutes of incubation, the first overnight, or after the second overnight step. We also tested samples taken after both overnight steps by mixing them with 100 µL of (z-Gly-Gly-Arg-7-amido-4-methylcoumarin (AMC).HCl substrate (Bachem 4002155.0025, Product #50-260-284, Fisher Scientific)). Fluorescence intensity was measured in technical duplicates (excitation at 370 nm, emission at 440 nm) every minute for 4 hours at 25°C. Commercial Streptokinase C (Product #S0577, Millipore Sigma) served as a positive control, and glu-PLG (Product #HCPG-0130, Prolytix) was used as a negative control. Fluorescence was measured on a SpectraMax M3, (Molecular Devices, San Jose, CA, USA), baseline corrected and means of the technical replicates for each experiment were plotted using Graphpad Prism.

### Protein stability predictors and surface accessibility calculation

The impact of individual amino acid substitutions was assessed using EvoEF2 (37) and FoldX (38), as described previously (31) using the structural information from the AlphaFold2-generated structure of SK_H46A_ (48). We calculated the relative solvent accessibility of each residue using the same structure and the DSSP v2.3 software (49). The raw solvent accessibility values were transformed by dividing each by the residue-specific maximum allowed solvent accessibilities in the empirical matrix provided in (50). The resulting relative solvent accessibilities were used for downstream analyses.

### Recombinant SK_H46A_ expression and purification

DNA coding for wildtype, M369F, K76L, G50L, and Y252T SK_H46A_ with a C-terminal His_6_-tag and TEV protease cleavage site were purchased from Twist Bioscience in a pET-24(+) expression vector carrying a kanamycin resistance gene. Plasmids were transformed into chemically competent NiCo21 DE3 *E. coli* (Product #C2529H, New England Biolabs). Protein expression and purification were performed as described in (51) .The cultures were grown in LB media at 30°C in log phase with shaking (OD600 0.4-0.6). IPTG (0.4 mM) was added for 2 hours followed by centrifugation of samples at 6000g x 20 mins at 4°C. The pellet was resuspended with 0.85% NaCl and the samples were spun again at 6000g x 20 mins at 4°C.The pellets were resuspended with buffer (25mM Hepes and 150Mm NaCl; (1X HBS pH 7.4)) and treated with 6.25 μM MgCl_2_, 0.6uM CaCl_2_, DNase (2mg/ml) and incubated with rotation at room temperature. The samples were freeze thawed in liquid nitrogen (1 min) and 37°C (5 min) three times before additional DNase (2mg/ml) was added. Finally, the volume was brought up to 5 mL to centrifuge at 16,000g x 20 mins. The resulting supernatant and pellet were checked for the presence of protein on a Coomassie gel, and the supernatant was purified by Fast Protein Liquid Chromatography (FPLC, AKTA Pure, Cytiva Corp, MA, USA). The SK protein was purified with a HiTrap TALON crude 1ml column (Cytiva) and equilibrated with 25mM HEPES containing 150mM NaCl and 5mM imidazole. The protein was eluted in 25mM HEPES containing 150mM NaCl and150mM imidazole and subsequently buffer exchanged into 25mM HEPES containing 150mM NaCl using a HiPrep 26/10 desalting column (Cytiva). The eluted protein fractions were pooled and stored at -80 °C and the purity of the protein assessed by SDS-PAGE followed by Coomassie staining.

The protein was then treated with TEV protease (Product # P8112S, New England Biolabs) according to the manufacturer’s instructions. To perform our assays, recombinant SK (15 μg) was treated with 1X TEV Protease buffer with or without TEV protease enzyme (reaction and control, respectively) at 4°C overnight. The samples were buffer exchanged with 1XHBS in 10KDa ultracentrifugal filters (Product# UFC901008, Millipore Sigma) to remove DTT from the TEV protease buffer and quantified spectrophotometrically (assuming a 1% extinction coefficient of 10).

### Mutational acceptance score calculations

Mutational acceptance scores for each position in SK were determined by averaging the log_2_-fold functional stability scores from our binding screen (27, 31). Using GraphPad Prism, we then applied a LOWESS regression with a 20-point smoothing window to the data. Areas with higher binding were defined as regions where the LOWESS curve was above the 75^th^ percentile value, while regions of lower binding were identified where the curve was below the 25^th^ percentile. These data were further used to generate heatmaps.

### Modified Halo/HoFF test assay

The Halo assays were performed based on previous reports (39, 52). Human α-thrombin (Product #HCATIII-0120, Prolytix, Essex Junction, VT, USA) was diluted to 240 nM in halo assay buffer (25 mM HEPES, 137 mM NaCl, 400 mM CaCl_2_, pH 7.4). Five microliters of this solution were added to each well of a white-walled, tissue culture-treated 96-well plate (Product#3610, Corning, Corning, NY), positioning the droplet against the wall. Human pooled plasma (25 µL; Innovative Research, Novi, MI), spiked with Alexa Fluor™ 488 Conjugate Fibrinogen (19.2 µg/mL; Product # F13191, ThermoFisher), and was dispensed into the thrombin droplet by circling the pipette tip along the bottom outer edge of the well, forming a halo-shape. The plasma mixtures were incubated for 1 hour at 37 °C to allow clot formation. Reaction mixes containing 25 mM HEPES, 137 mM NaCl, 2.5 mM Ca^2+^, 0.8 mM S-2251 (plasmin chromogenic substrate; DiaPharma, West Chester, OH, USA), and 12.8 nM of a given SK variant were prepared for triplicate wells (final well concentrations: 8 nM SK, 0.5 mM S-2251). The plate reader was preheated to 37 °C. After warming, 50 µL of reaction mixture was added to each well, and absorbance at 405 nm (S-2251 cleavage) and fluorescence (excitation 488 nm, emission 530 nm; Alexa488-fibrin cleavage) were measured every 90 seconds for 1 hour and 20 minutes at 37 °C. The absorbance and the fluorescence were measured simultaneously on a BioTek CYTATION5 imaging reader (BioTek, Winooski, VT, USA). Absorbance data were baseline-corrected, and the means of the three replicates were plotted as a function of time for each variant.

## Data analysis and Code

All code for data analysis has been deposited on GitHub (https://github.com/SrishtiBaid/SK_PLG_binding.git), and HTS sequencing results are available at https://doi.org/10.7302/d3aw-z910.

## Acknowledgements

Funding was provided by AHA Postdoctoral Fellowship: 25POST1372745 (SB), and NIH R35HL171421 (DG). UM-GPT was used to refine the language and the manuscript flow. The authors thank Khushali Kataria for assistance with the schematic figures of this manuscript.

