## Supplementary data for "Mapping the Mutational Landscape for Streptokinase Binding to Plasminogen"

Supplementary table 1: Primer table

| | |
|---|---|
| SK cloning seq check FP | TTCATGCTGCCGGCTTTCTCGG |
| SK cloning seq check RP | TCGTCATCGTCCTTGTAGTCACCACCA |
| SK Error Prone FP<br>AscI-6XHis-TEV-SK-FP | AGCCGGCCGGCGCGCCACACCATCACCATCACCATGAAAACCTGTATTTTCAG |
| TEV intocodingSK FP | CACCATCACCATGAAAACCTGTATTTTCAG |
| SK Error Prone PCR RP<br>IntocodingSK-NotI RP | CGTCATCGTCCTTGTAGTCACCACCA |
| Amplicon Seq Primer 1 FP | CACCATCACCATGAAAACCTG |
| Amplicon Seq Primer 1 RP | GATTTCAAAAAATTTAAGACTAATGTCTTGA |
| Amplicon Seq Primer 2 FP | GGTACTGTTGAGGGGACG |
| Amplicon Seq Primer 2 RP | TTTATGTGACATCGCGCC |
| Amplicon Seq Primer 3 FP | AAATCAAAACCATTTGCTACTGA |
| Amplicon Seq Primer 3 RP | GGTTGCATCGCTTGCAAAAT |
| Amplicon Seq Primer 4 FP | CGACGACTACTTTGAGGTC |
| Amplicon Seq Primer 4 RP | CTAACGCGCACATGTCC |
| Amplicon Seq Primer 5 FP | CCTGTCCAAGAATTTTTGCTAAG |
| Amplicon Seq Primer 5 RP | TTTGAGACCTGGTCTGAAATC |
| Amplicon Seq Primer 6 FP | AGTTTACTCCCTTAAACCCTGA |
| Amplicon Seq Primer 6 RP | TATAGCCTGGGTGGTTTTTG |
| Amplicon Seq Primer 7 FP | GCTCAAGCACAAAGCATT |
| Amplicon Seq Primer 7 RP | GCTCAAGCACAAAGCATT |
| Amplicon Seq Primer 8 FP | TTTACCAATGGATCAAGAGTTTAC |
| Amplicon Seq Primer 8 RP | TACGGCTTTTCCCCTTTTTT |
| Amplicon Seq Primer 9 FP | CTGATCTCTGAGAAATATTACGTCC |
| Amplicon Seq Primer 9 RP | TTCGCTAGCTGTTAAGAGC |
| Amplicon Seq Primer 10 FP | ACGAATTGCTAAAAAGTGAGC |
| Amplicon Seq Primer 10 RP | CTACTTTTCCAGTTAAGGTATAGTCC |
| Amplicon Seq Primer 11 FP | CAATCTCGATGCTTTTGGTATT |
| Amplicon Seq Primer 11 RP | CGATCTTTATCATAGGCTAAATGAT |
| Amplicon Seq Primer 12 FP | GCGACCCGAAGGAGAGAATG |
| Amplicon Seq Primer 12 RP | GCCGCCGCTGAACCA |

Supplementary Figure 1

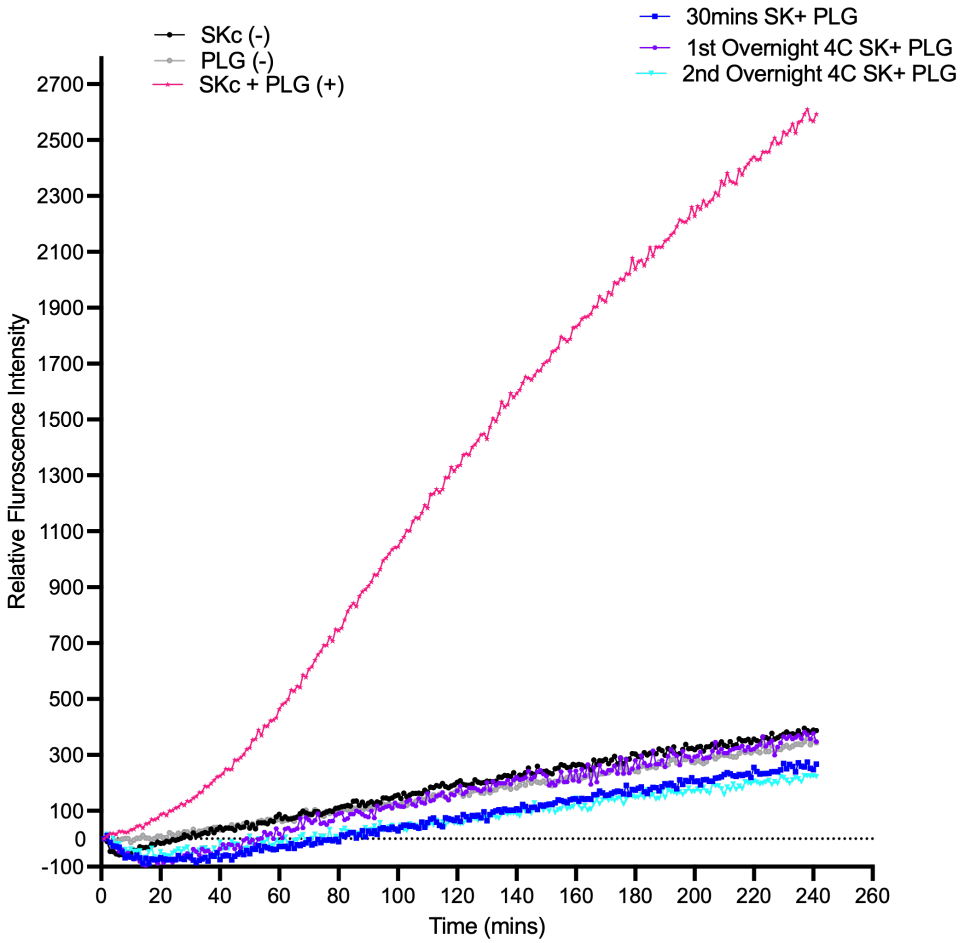


**Supplementary Figure 1**: **Activation of PLG to $PLG_{SK}$ leads to the rapid generation of plasmin.** Plasmin has a higher affinity for SK than does PLG (1, 2). To determine whether any functionally active $PLG_{SK}$ is being generated in our assay, due to the possibility of residual SK activity, despite a protected $Ile^1$ , as suggested by a previous report (3). No background activation was detected in the SK and PLG interaction experiments. We measured fluorescence over 4 hours at various stages of the experiment- including after 30 minutes at room temperature and after two overnight incubations at 4°C. Commercial Streptokinase C (SKc) served as the positive control (showing PLG binding, pink line), while samples with only SKc or only PLG served as negative controls. Fluorescence readings were baseline-corrected, and the average values from duplicate experiments (n=2) were plotted over time for each variant in Graphpad PRISM.

Supplementary Figure 2

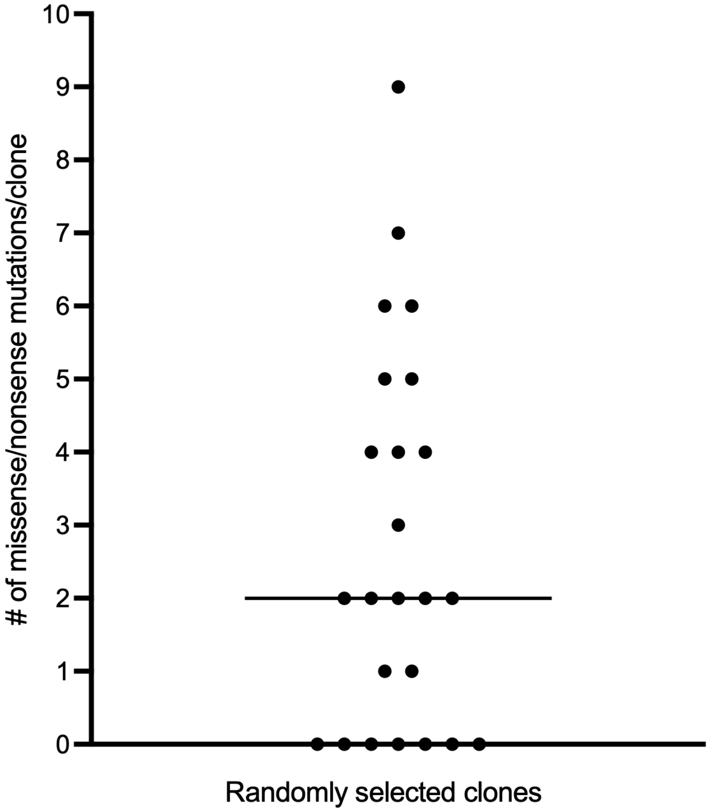


**Supplementary Figure 2: Sanger sequence of the error prone PCR library identified a median of 2 mutations per clone.** The number of missense/nonsense mutations that were identified in each of 24 randomly selected phage clones from Sanger sequence are shown here.

Supplementary Figure 3

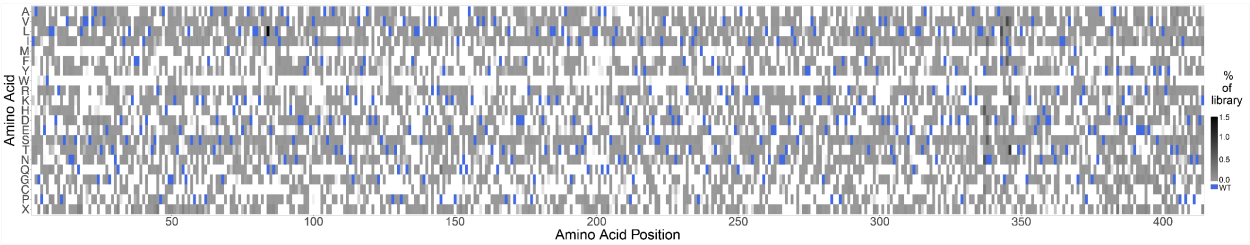


**Supplementary Figure 3**: **Library coverage for recombinant SK**. The error prone PCR library for SK utilized in this study encompasses 71% of all possible single amino acid substitutions. Amino acid positions within SK are displayed along the x-axis, and the potential amino acid substitutions (X represents nonsense mutations) are plotted on the y-axis. The relative abundance of each variant in the input library, as measured by Illumina sequencing, is depicted using a greyscale gradient, with the wild type residues at each position highlighted in blue. White indicates no coverage at that position.

Supplementary Figure 4

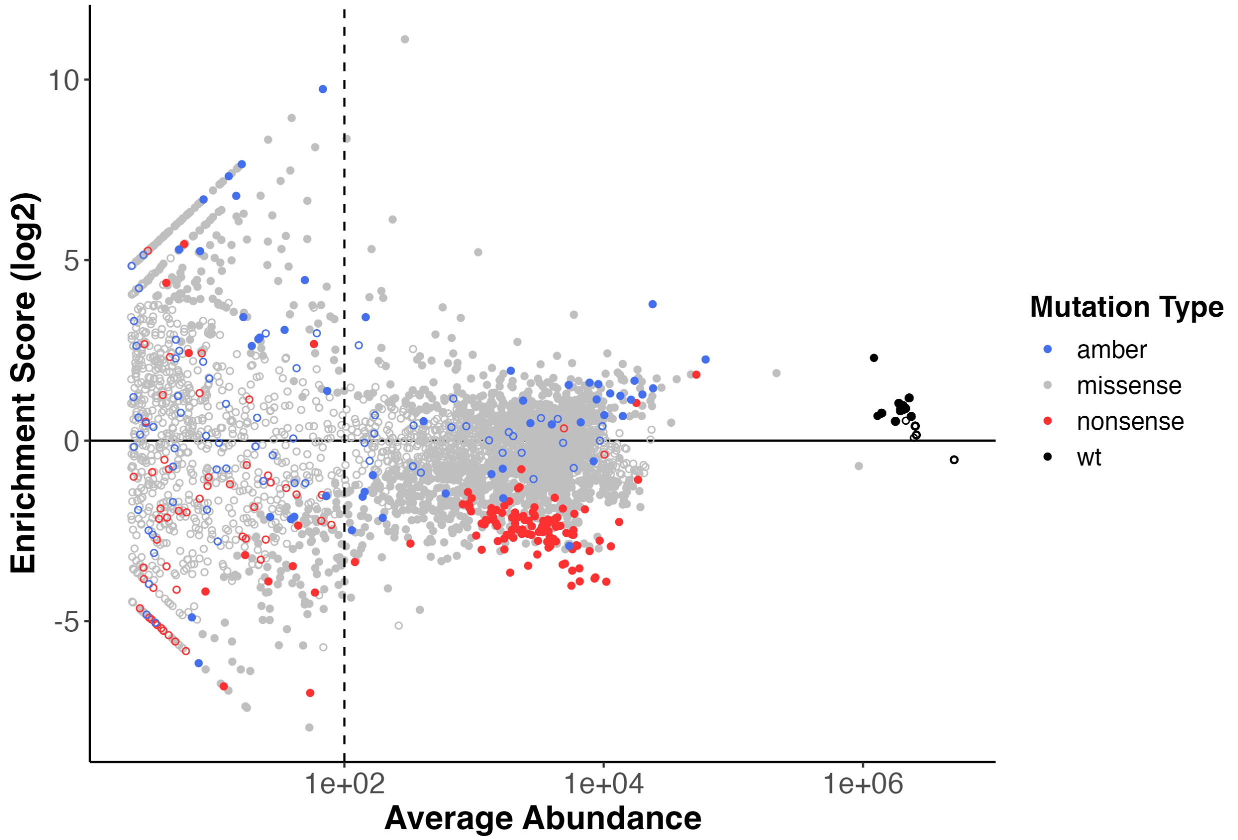


**Supplementary figure 4**: **MA plot showing distribution of mutations.** PLG- SK selected libraries contained variants that either lost or gained binding function. The MA plot (4) shows the average abundance vs $log_2$ fold change for individual mutations. Wildtype are indicated in black, missense in grey, and amber stop in blue and the other nonsense mutations in red. A base mean or average abundance score of 100 was set as a cut-off to identify threshold for significance. The average abundance is defined as average counts for the input and selected library. The variants with closed circles correspond to $p_{adj}$ <0.01, classified as significant (with open circles not significant). The figure was generated using the ggplot2 package in R studio (5).

Supplementary Figure 5

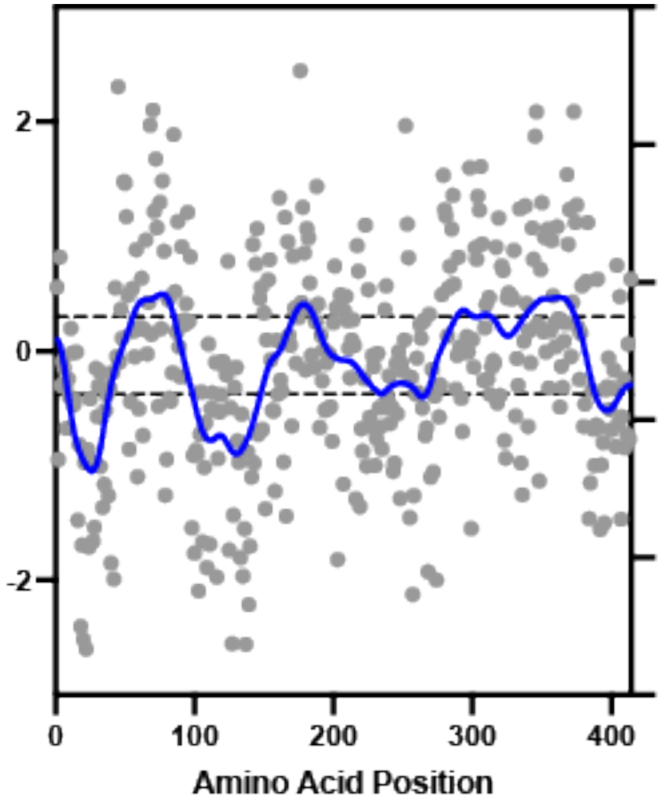


**Supplementary figure 5**: **Calculated mutational acceptance scores**. The log2-fold change results from the binding screen, corrected for SK variants shown in the heat map, were summed at each amino acid position to calculate mutational acceptance scores (shown in grey). These scores were then plotted against amino acid position and fitted with a LOWESS regression (blue line). Dotted lines mark both the 25$^{th}$ and the 75$^{th}$ percentile of the LOWESS regression curve.

Supplementary Figure 6

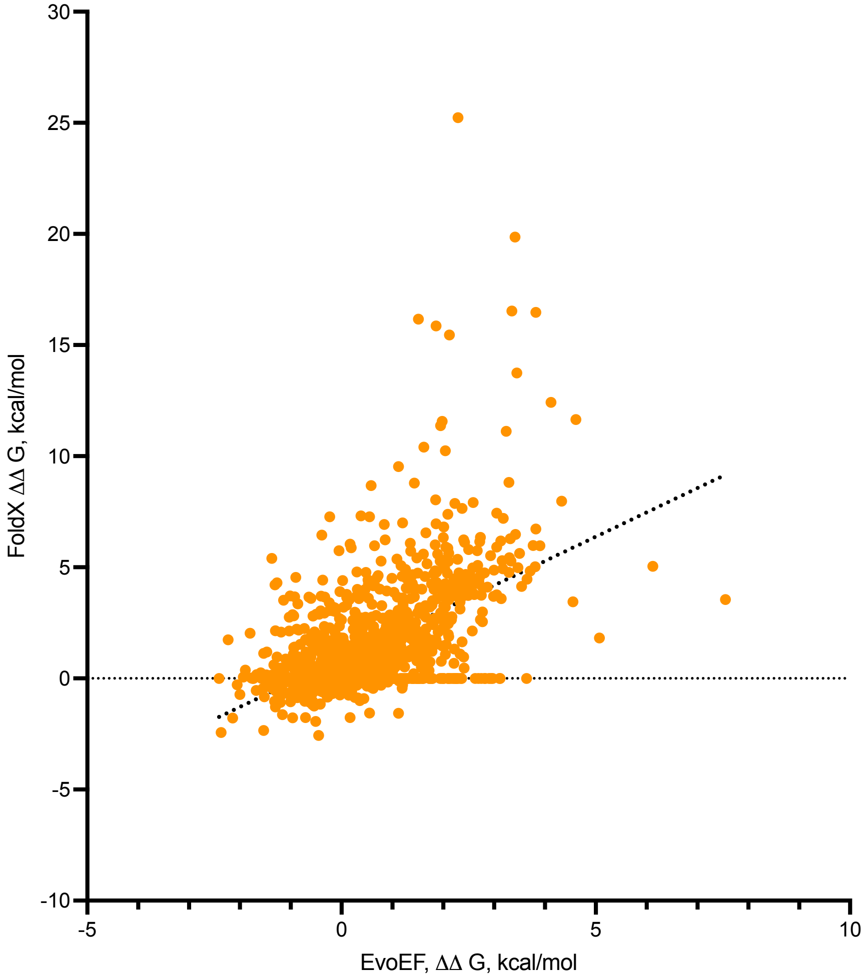


**Supplementary figure 6**: **Comparison of $\Delta\Delta G$ values for SK amino acid substitution using EvoEF and FoldX**. $\Delta\Delta G$ values for SK amino acid substitution was calculated using 2 stability predictor computational models, EvoEF and FoldX, by analyzing the effects of amino acid substitutions in SK. The correlation between the 2 models is indicated for each amino acid substitution (orange), with the regression plotted. The two models show a significant correlation (y-intercept = 0.9031, slope = 1.095, R2 = 0.2930, p < <0.0001).
